# Simpler predictive models provide higher accuracy for ovarian cancer detection

**DOI:** 10.1101/2024.12.19.629553

**Authors:** Derrick E. Wood, Joseph Roy, Bari J. Ballew

## Abstract

Ovarian cancer remains a danger to women’s health, and accurate screening tests would likely increase survival. Two established protein biomarkers, CA125 and HE4, have been shown to work well in isolation, but achieve even higher accuracy when combined using logistic regression (LR). We show here that this LR-based combination of protein concentrations achieves high accuracy when distinguishing healthy samples from cancer samples (AUC = 0.99) and benign masses from cancer (AUC = 0.86). This approach exhibits superior performance on an external validation cohort compared to a more complex method, which was published with the dataset we use here and tested on the same data. While our method only uses proteins, the more complex method also uses features derived from cell-free DNA (cfDNA). We show that many of that method’s cfDNA features are affected by confounding technical variation, which impacts the previously reported results. Our results are in line with the principle that simpler machine learning models will tend to exhibit better generalizability on new data.

## Introduction

In 2024, the American Cancer Society projected 19,680 new cases of ovarian cancer in the United States, with an estimated 12,740 deaths attributed to the disease [1]. The prognosis for ovarian cancer is significantly poorer when diagnosed at advanced stages compared to earlier detection [2]. Therefore, the development of accurate assays for early ovarian cancer detection is crucial for improving survival rates. Such assays could be employed in two key scenarios: as a screening tool for asymptomatic individuals, and as a diagnostic aid for patients with an ovarian mass to assess its malignancy.

Assays for cancer detection can use one or more biomarkers. Genomic biomarkers such as somatic mutations in the BRAF gene [3], homologous recombination repair deficiency, and mismatch repair deficiency [4] are frequently analyzed for ovarian cancer diagnosis and treatment. The proteins CA125 and HE4 also serve as common protein biomarkers for diagnosing ovarian cancer [2]. Elevated levels of both CA125 and HE4 are linked to ovarian cancer [5,6], though HE4 concentrations also rise with age [7].

Researchers have developed algorithms that combine these protein biomarkers to enhance diagnostic accuracy. For example, in 2009, researchers introduced the Risk of Ovarian Malignancy Algorithm (ROMA), a logistic regression (LR) model that integrates CA125 and HE4 values, achieving superior diagnostic precision compared to either protein individually [8]. ROMA selects its LR model based on a patient’s menopausal status. In contrast, the Copenhagen Index [9] utilizes a patient’s age along with CA125 and HE4 serum levels in an LR model to estimate the probability of cancer.

Other cancer detection approaches have targeted both screening and diagnostic applications. DELFI [10,11] utilizes an algorithm to detect cancer by analyzing fragmentation patterns in cell-free DNA (cfDNA) sequences, alongside copy number information. More recently, we have seen the introduction of DELFI-Pro [12], a computational pipeline that employs logistic regression (LR) to integrate CA125 and HE4 measurements with numerous cfDNA-derived features from DELFI. For ovarian cancer detection, DELFI-Pro demonstrated superior performance compared to both CA125 and HE4 protein concentration levels. The authors also noted that these two proteins had the largest magnitude coefficients in their screening model, collectively accounting for 68.8% of the total feature importance, while features associated with copy number changes contributed 24.1%.

In their experimental design for the DELFI-Pro article, the authors noted that they took measures “to reduce the possibility that differences between patients with or without cancer were not due to batch variability.” Batch effects, a well-recognized issue in laboratory analyses [13], represent technical variations that correlate with the processing batches of samples, rather than underlying biological or scientific factors [14,15]. These confounding effects can significantly impact downstream analyses, potentially leading to overly optimistic results, even when cross-validation is employed [16,17]. Notably, batch effects are known to affect various aspects of nucleic acid processing projects, including GC-content bias [18], RNA transcript expression [19], and copy number [20,21].

Related to concerns of including batch-affected features, an important consideration in using machine learning models such as LR is to ensure that the size of the feature set is minimal while still providing adequate performance [22]. As others recently stated [23]: “A model that has good accuracy and low complexity, *i.e.* a model with fewer input features, will be much more robust to perturbations, faster and easier to understand, and much cheaper to maintain and update.” Our own desire to follow the principle of parsimony motivated us to examine the DELFI-Pro data and determine how well LR models using only the measurements of CA125 and HE4 would perform. Such models, with similar architecture to that of the long-established ROMA, might give firmer insight into the added predictive power of DELFI-Pro’s cfDNA-derived features. We present the results of this examination here.

## Results

### Collection of protein concentration data

Briefly, the blood samples from which the DELFI-Pro authors obtained their data were from two cohorts: a “discovery” cohort of samples from European studies, and a “validation” cohort of samples from United States studies. Protein concentration levels for CA125 and HE4 were quantified using immunoassay measurements from these blood samples, in most cases from plasma, although a minority of measurements were made from serum samples.

In both the discovery and validation cohorts, samples were collected from three classes of donors: those with no known ovarian lesions (labeled “healthy” hereafter), those with benign adnexal masses, and those with ovarian cancer. Both cohorts have two overlapping datasets: a “screening dataset” consisting of the healthy samples and the cancer samples; and a “diagnostic dataset” consisting of the samples from benign masses and the cancer samples. The two datasets allow building of classification models in both a screening and a diagnostic context, and evaluation of the predictive power of an approach in both contexts. To facilitate comparison between the models we built and the DELFI-Pro models, we maintained this same division of cohorts and use of classification contexts.

### Building a protein-only logistic regression model

We chose to use a logistic regression model architecture for our machine learning approach. Other models are of course possible, but bearing in mind that our input data was only two-dimensional, that ROMA had achieved good performance with such an LR model, and that DELFI-Pro also utilized LR, we felt LR was the appropriate choice here. To mitigate overfitting, our LR implementation incorporated a regularization penalty, a technique also employed by DELFI-Pro. Our initial analysis focused on the discovery cohort to establish parameters for the LR model and determine appropriate data preprocessing steps.

A common and important early step in machine learning with laboratory analytes is to use the zlog transformation [24]. This transformation compensates for the fact that the range of values from pathological samples can be an order of magnitude higher than that seen in healthy samples. The zlog transformation takes the logarithm of the input value so that the range of data is reduced, and also standardizes the result so that healthy data tends to be close to a value of zero. We applied this transformation to CA125 and HE4 values, using the zlog values for subsequent analyses instead of the original protein concentrations. In this zlog feature space, we can see that CA125 appears to distinguish between healthy and cancer samples well, but that HE4 may be needed to appropriately distinguish between benign masses and cancer samples (Supplementary Figure S1). The two zlog features are not strongly correlated, with a Pearson’s R of 0.57 in the discovery cohort.

Similarly to the DELFI-Pro publication [12], we performed a 5-fold stratified cross-validation, repeated 10 times, to assess each model’s performance against its respective discovery data. We recorded the mean score of a sample across all 10 repeats of cross validation. To evaluate the performance of a model, we use the receiver operating characteristic (ROC) area under the curve (AUC) [25]. After cross validation, our screening protein-only model had an AUC of 0.96, equal to the 0.96 reported by DELFI-Pro (Figure 1). In the diagnostic context, our protein-only model had an AUC of 0.86, slightly less than the cross validation AUC of 0.88 reported by DELFI-Pro (Supplementary Figure S2).

**Figure 1.**
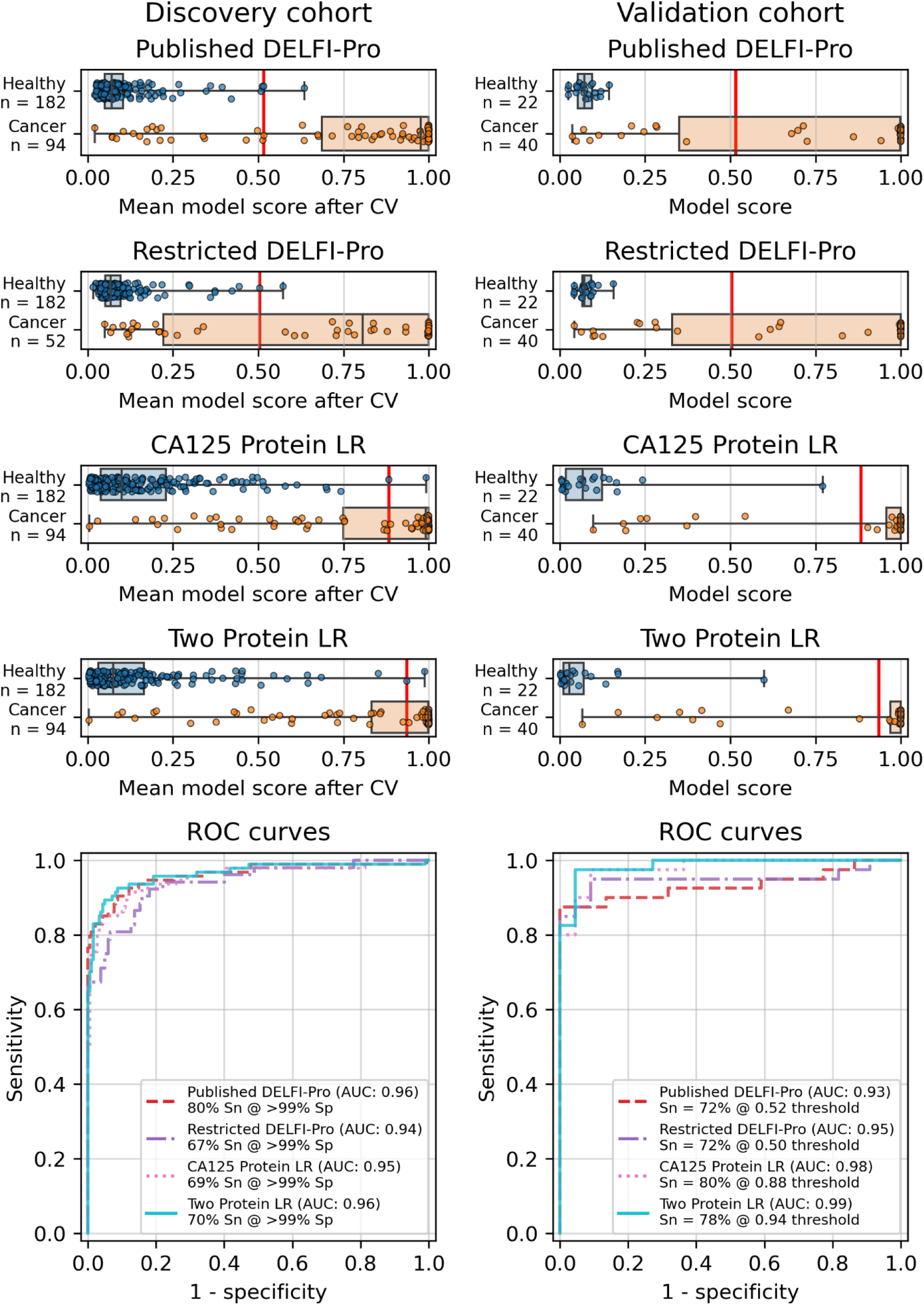
Logistic regression model performance in a screening context. Cross validation (“CV”) results for the discovery cohort, and validation results on the validation cohort are shown for the published DELFI-Pro screening model, a CA125 protein logistic regression model, and a two-protein logistic regression (“Two Protein LR”) screening model that combines CA125 and HE4. These models seek to distinguish between samples from healthy donors and samples from patients with ovarian cancer. We trained an additional DELFI-Pro model (“Restricted DELFI-Pro”) by removing from the discovery cohort the 42 cancer samples that had a “PGDX” prefix. Receiver operating characteristic (ROC) curves demonstrate the performance of the two models at various score thresholds, with the area under the curve (AUC) growing proportionally to model accuracy. Model sensitivity (“Sn”) is reported at the clinically-relevant >99% specificity (“Sp”) threshold using the discovery cohort CV results, and the corresponding score threshold is marked as a vertical red line on the score plots in the top half of the figure.

In addition to this cross validation work, we also fit the same models to the full discovery datasets, so that the models could be examined and then run on the validation cohorts. Examining the slopes of the models’ decision boundaries (Supplementary Figure S3), we see that the screening model is influenced slightly more by CA125’s value than HE4’s, with a decision boundary slope of -1.23. In contrast, the diagnostic model’s boundary line is far more horizontal with a slope of -0.12, indicating that HE4 influences that model more. The screening model has a CA125 term coefficient of 1.25, an HE4 term coefficient of 1.02, and an intercept of -2.22. The diagnostic model has CA125 and HE4 coefficients of 0.14 and 1.15, respectively, with an intercept of -1.66.

### Validation and performance of the fitted models

While cross validation results are useful, testing on a separate and unrelated dataset is considered to be a more reliable test of a model’s generalizability [26]. As did the DELFI-Pro authors, we ran the models fit to the discovery cohorts on the corresponding validation cohorts. While our protein-only models performed very similarly to DELFI-Pro in cross validation, our models substantially outperformed DELFI-Pro on the validation cohorts. Our screening model achieved an AUC of 0.99, above the corresponding DELFI-Pro AUC of 0.93 (Figure 1). Similarly, our diagnostic model achieved an AUC of 0.86, exceeding the DELFI-Pro diagnostic validation AUC of 0.81 (Supplementary Figure S2).

We also compared our screening model against results from another model created by the DELFI-Pro authors. We refer to this other model, which is a combination of only CA125 and HE4 using logistic regression, as “DELFI-noDNA” here due to its lack of cfDNA features. The DELFI-noDNA model’s cross validation results were published as a supplementary figure in the original publication [12], but the validation results are not present in that figure, nor do we find any mention of this model or its results discussed in that publication’s text or tables. Both our screening model and the DELFI-noDNA model had cross validation AUCs of 0.96 on the discovery cohort. In the validation cohort, our screening model’s AUC of 0.99 was slightly higher than the DELFI-noDNA model’s AUC of 0.98 (Supplementary Figure S4).

### Batch effects confounded copy number features

Because the DELFI-Pro models exhibited lower accuracy on validation data when compared to the discovery cohorts, we suspected a confounding factor may be affecting the cfDNA features. Reviewing the heatmap published with DELFI-Pro [12], we saw that several copy number-related features appeared to be: (a) close to zero in all benign and healthy samples; (b) negative in the vast majority of cancer samples; and (c) uncorrelated with the DELFI-Pro score in cancer samples. These facts appeared incongruous to us, as a feature that is negative only in cancer samples would intuitively be well-correlated with the probability of a sample belonging to a cancer patient.

After investigating the code and data used to generate the heatmap, we found that the final 85 samples in the dataset had sample IDs that shared the prefix “PGDX”, as opposed to the other 394 samples with IDs with a “CGPL” prefix. We then created our own heatmap, grouping samples by cancer status and sample prefix. Our heatmap, focused on copy number and protein features, shows that several copy number features have values that are lower in PGDX samples compared to the CGPL samples (Figure 2). These results are consistent with the conclusion we reached upon our reading of the DELFI-Pro heatmap code: because the data of the heatmap was not properly joined with the row labels, the reordering of the *row labels* did not cause the heatmap’s corresponding *data rows* to be reordered from the initial ordering within the input file. The connection we saw between negative copy number features and cancer samples in the original publication, then, appears to be merely an artifact of batch effects and an unfortunate error in data visualization.

**Figure 2.**
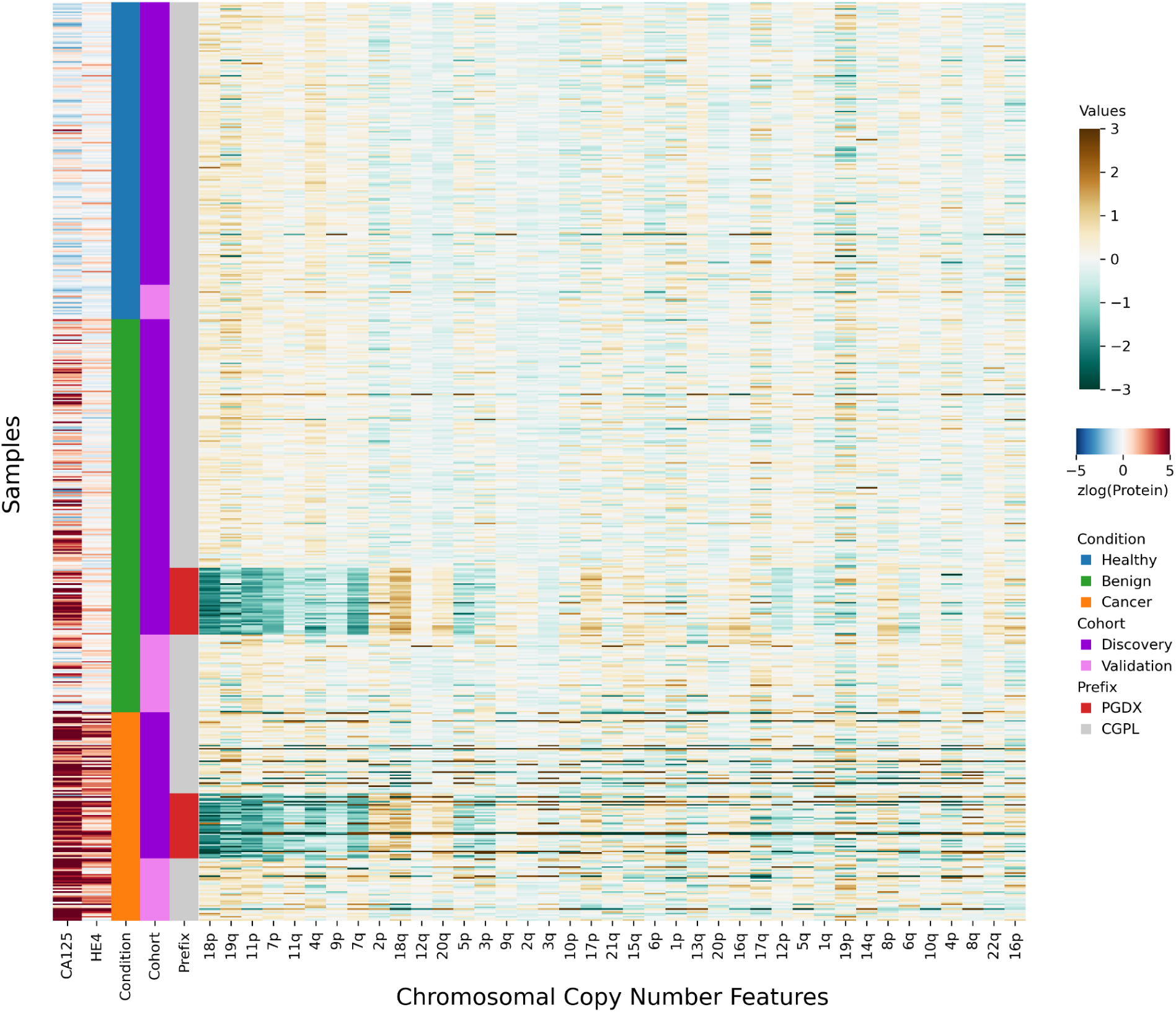
Heatmap demonstrating sample prefix-associated effect on copy number features. The samples from the DELFI-Pro dataset are shown here, with each row representing features from a single sample. For the proteins CA125 and HE4, we display the zlog-transformed values for each sample. Features are sorted such that the leftmost features are more affected by the batch effect. All features had values standardized here prior to display.

Seeing the visually apparent differences in PGDX and CGPL samples within the copy number features, we sought to quantify the predictive value of the features absent this confounding factor, as well as the impact of the differences. Because the CGPL samples were 4.6 times more numerous than the PGDX samples, and the CGPL samples included all three of our conditions (healthy, benign, and cancer), we focused our analysis first on the CGPL samples.

Within the CGPL samples of the discovery cohort, we calculated the AUC for cancer prediction of each copy number feature as well as our two protein features (Supplementary Figure S5). In a screening context, the AUCs of the two protein features were both above 0.9, indicating strong association with cancer for each individually. In a diagnostic context, HE4 had an AUC of 0.90, and CA125 had an AUC of 0.79, results which are congruent with the LR coefficients we reported earlier. None of the 39 copy number features had an AUC above 0.75 in either the screening or diagnostic contexts. This indicates that within the CGPL samples, copy number features were individually not strongly associated with cancer.

Next, we examined only the cancer samples within the discovery cohort (52 CGPL and 42 PGDX samples), calculating the AUC for predicting the sample prefix (Supplementary Figure S5). In this experiment, neither protein had an AUC of more than 0.56. However, 44% (17 of 39) of the copy number features had AUCs above 0.75, with three (corresponding to changes to chromosome arms 11p, 18p, and 19q) having AUCs above 0.9.

All samples labelled with a PGDX prefix in the repository metadata were listed with CGPL prefixes in the original paper’s supplementary data, but we are able to match samples across the two namespaces by comparing protein concentration values. Reviewing the metadata associated with the PGDX samples showed that all PGDX samples were processed as part of library batches 1-16, and all CGPL samples were processed as part of library batches 17-38. We also see that the PGDX samples contain no healthy or validation samples (Figure 2, Supplementary Figure S6). We are unable to determine from the original publication what caused this cohort to be processed with a different sample prefix.

To further explore the impact of the observed technical confounding, we conducted an experiment suggested by some of the DELFI-Pro authors during this article’s review. This involved creating a “restricted DELFI-Pro” screening model by removing PGDX samples from the discovery cohort and training the new model on this reduced cohort. Our methodology, including cross-validation on the discovery cohort and external validation on the validation cohort, remained consistent with the original DELFI-Pro publication [12], with the sole exception of the exclusion of 42 cancer samples with a PGDX prefix.

The results showed a decrease in AUC during cross validation, from 0.96 in the original DELFI-Pro model to 0.94 in the new restricted model. Conversely, in the validation cohort, the AUC increased from 0.93 to 0.95 (Figure 1). We also observe significant shifts in feature importance. For example, chromosome 18q, which was the second most important chromosomal change in the published DELFI-Pro screening model, now has a zero coefficient in the new model. In contrast, chromosome 10p, which previously had no impact, now exhibits the most influential Z-score in the new model. We present a detailed comparison of feature importances between the two models in Figure 3.

**Figure 3.**
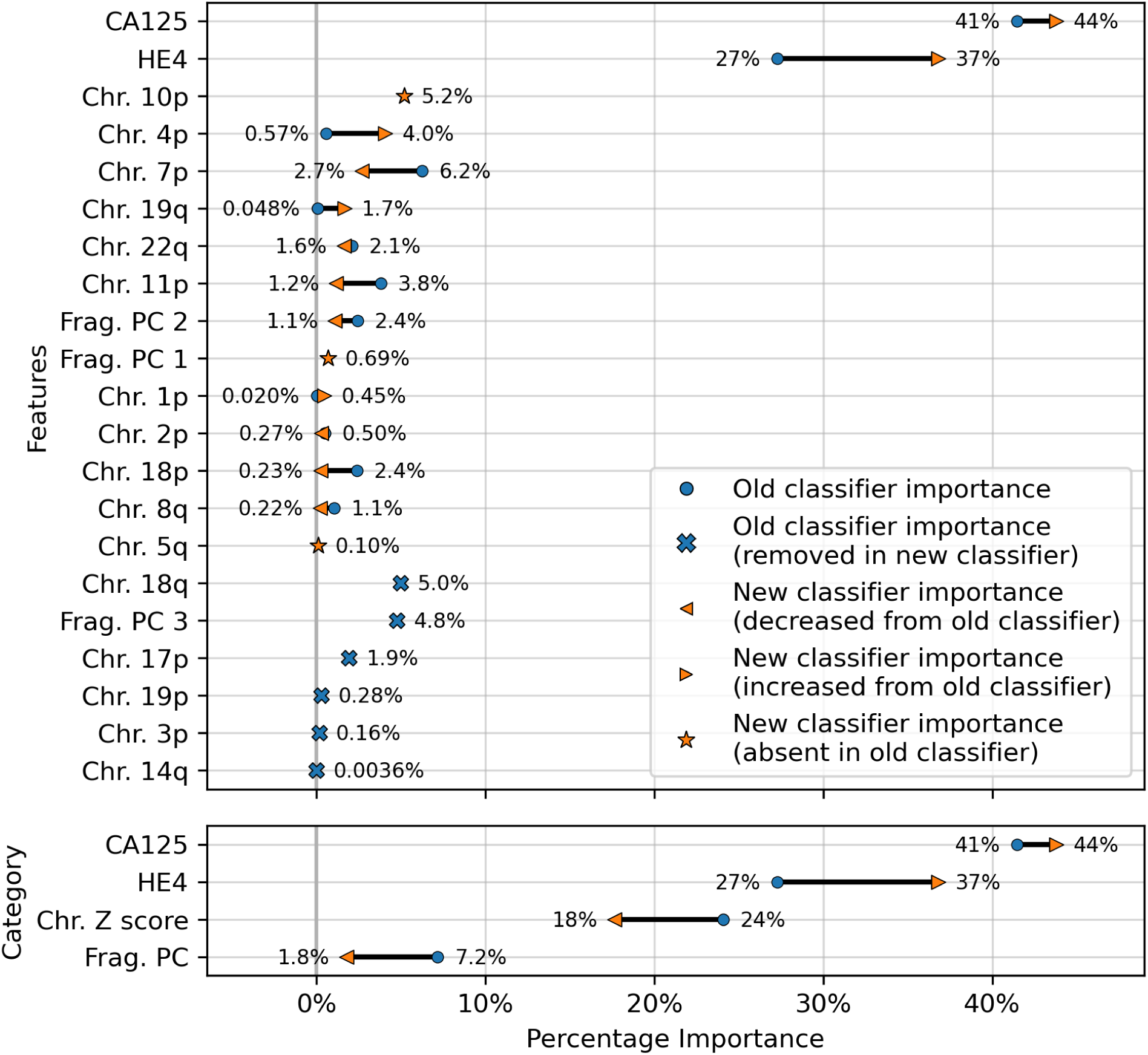
Comparison of feature importances for two DELFI-Pro screening models. Using the same model architecture as the DELFI-Pro authors, we built a screening model trained on the discovery cohort without the PGDX samples. This new classification model was compared to the old model (the published DELFI-Pro screening model) above, and feature importances were calculated. The top panel shows the differences in feature importance for individual features, and the bottom panel shows the differences in importance for the same 4 categories of features reported by the DELFI-Pro authors. The categories include the two proteins CA125 and HE4, along with chromosomal copy number Z scores and fragmentation principal component (“PC”) values. Any feature not present in the figure had a coefficient of zero in both models.

Finally, an important matter is clinically relevant specificity in screening. The DELFI-Pro authors rightly note [12] that extremely high specificity is necessary to ensure a high positive predictive value (PPV) for a screening test, given the low prevalence of ovarian cancer. Such high PPV is necessary to minimize false positives and the unnecessary concern and procedures that would follow from those false positives. Therefore, in evaluating the new restricted DELFI-Pro model, we calculated the LR model score threshold that would correspond to >99% specificity within the cross validation data. We report the sensitivity of each model at that model’s threshold in Figure 1. Notably, this sensitivity at >99% specificity for the restricted DELFI-Pro classification model is lower than that of the published DELFI-Pro model (67% vs. 80%), which is consistent with our expectations regarding the impact of the technical confounding on the published DELFI-Pro results.

Because CA125 has been used in other ovarian cancer screening trials [27–29], we also report CA125’s performance as a screening tool here, using logistic regression to determine an appropriate threshold. We performed the cross validation and validation testing in a similar manner to our tests of the two-protein model. At the >99% specificity threshold, CA125 has a cross validation sensitivity of 69%, slightly less than the 70% of our two-protein screening model (Figure 1). With 182 healthy samples in the discovery cohort, the point estimate for specificity that clears the >99% threshold is 99.45% (181/182). Given the 0.145% prevalence in the United States for ovarian cancer that SEER reported in 2022 [30], our two-protein model’s PPV would be 15.6% at this high specificity threshold. The estimated PPVs for the CA125 and restricted DELFI-Pro models are very similar to that of our two-protein model, at 15.4% and 15.1%, respectively. The CA125 model’s 99.45% specificity threshold of 0.88 corresponds to a zlog value of 2.78, or 52.2 U/mL. The CA125 model has a single coefficient of 1.39, with an intercept term of -1.84.

## Discussion

Our findings largely align with those of the DELFI-Pro authors regarding the predictive utility of the proteins studied. While CA125 and HE4 concentrations are individually valuable biomarkers, their predictive power increases when combined within a machine learning model. We extended the analysis beyond that of the DELFI-Pro publication to specifically evaluate and thoroughly report the performance of these two proteins independently of any cfDNA-associated features. We also discovered a strong batch effect within the data used to train the DELFI-Pro models, and analyzed its impact on the accuracy of the published DELFI-Pro screening model.

When compared to the DELFI-Pro models’ performance on the validation cohorts, we see superior performance from our simpler models. We also see concordant performance with a protein-only model with full results available in the DELFI-Pro code repository, giving us further confidence in the results of our parsimonious approach. Given these validation results and the very similar cross validation performance between our models and DELFI-Pro on the discovery cohorts, it appears that the simpler models exhibit improved generalizability compared to the DELFI-Pro models. Our protein-only models would also tend to be easier, faster, and cheaper to perform than the DELFI-Pro approach. This is because a laboratory test using our models would only require the quantification of protein concentrations, rather than also needing to prepare, sequence, and bioinformatically analyze DNA libraries. In light of these results, the additional predictive utility over CA125 and HE4 that would be provided by DELFI-Pro’s cfDNA-derived features remains unclear.

Our analysis revealed a substantial batch effect in the discovery cohort, which negatively impacted the DELFI-Pro models’ performance on validation data. This decline is likely due to the models incorrectly correlating technical variations with clinical labels, a consequence of the dataset’s construction. The absence of healthy samples in the affected batches prevented the screening model from recognizing that technical artifacts, present only in cancer samples, were indeed just artifacts.

While later batches generally contained a mix of conditions, the initial 16 batches (those prefixed with “PGDX") were largely homogenous. Only batch 10 included both cancer and benign samples (Supplementary Figure S6). This lack of condition diversity in the early batches, combined with pronounced effects on copy number features, can lead to inaccurate models. Consequently, the importance of copy number features, as reported by the DELFI-Pro authors, is likely attributable to these batch effects rather than biological factors (Figure 3, Supplementary Figure S5).

We do not report evidence of strong batch effects within any of the validation data, but we note that the mixing of validation and discovery data within batches (Supplementary Figure S6) constitutes a form of data leakage [31]. Such data leakage could result in overly optimistic validation results, because machine learning models could effectively learn batch-specific information that would not occur in future batches. To prevent this, we recommend that validation samples are both collected and processed after any samples used to train models. This creates a time separation between the two cohorts that can provide a much stronger assurance that, when the model is deployed, validation results will generalize to clinical performance.

Although our simpler models appear to be more generalizable than the DELFI-Pro models, these results do not necessarily mean that any of our protein-only models will perform as well on a new validation dataset as they did on the validation cohorts we analyzed here. Our models’ results are necessarily limited to the DELFI-Pro dataset, and the workflow used to produce that dataset. A more comprehensive validation would be necessary to show that a specific model, paired with a specific laboratory workflow, would perform well enough to be used as a clinical screening or diagnostic tool. Such a validation should detail a consistent biofluid from which the measurements will be taken, as opposed to the combination of plasma and serum that we find in the DELFI-Pro experiments. The training data for this more substantive validation should also be composed of heterogeneous batches, with the proportions of the various conditions being similar from batch to batch. And finally, as we previously recommended, all validation samples would need to be processed subsequent to both the laboratory and analytical processing of the training data.

We also note that even though our results suggest the two-protein screening model outperforms CA125 alone, the difference in performance is quite small. The cost of analyzing both CA125 and HE4 may not be justified for ovarian cancer screening given the small gain in performance of the two-protein model. Further work would be needed to establish whether the combination of proteins provides sufficient additional accuracy for HE4 to be added into a clinical screening test.

The presence of confounding batch effects on the DELFI-Pro copy number features points to an important concern in clinical diagnostics. Experimental design to mitigate the effects of batch variation is essential, but this must also be paired with robust quality control and data analyses to either correct for batch variation or disqualify samples when extraordinary variation is encountered. The absence of such checks can lead to the creation of screening or diagnostic models that do not generalize to new samples. Such a lack of generalizability in turn can lead to a costly failure to pass a well-designed validation experiment, or worse, elevated rates of inaccurate clinical results if the validation experiment is not well-designed.

While cfDNA-based diagnostics are certainly not unsuitable for clinical tests, their additional benefits over other approaches must be clearly demonstrated for specific applications. Beyond the cost savings mentioned earlier, a method solely examining protein concentrations would necessitate fewer and less expensive quality control checks than one incorporating DNA sequencing. Throughout our comparisons and analysis, we had hoped to demonstrate these additional benefits of DELFI-Pro’s cfDNA features directly. We do note the apparent visual concordance between copy number results in ovarian cancer tumors in TCGA [32] and the DELFI-Pro copy number features, as presented by the DELFI-Pro authors [12]. However, our results with the restricted DELFI-Pro screening model and the lower importance of copy number features seen in Figure 3 make these features’ additional value uncertain. They may be a group of informative features for ovarian cancer detection, but we do not have evidence that they provide value beyond that provided by the two proteins studied here.

It is also worth noting that while both we and the DELFI-Pro authors report sensitivity at a clinically relevant >99% specificity, our methods for calculating such a threshold differ in important ways. We selected the minimal threshold that would be at least 99% specific in the discovery cohort, which is equivalent to 99.45% specificity. The DELFI-Pro authors report a threshold that is above the highest-scoring healthy sample in the discovery cohort, corresponding to 100% specificity (“no false positives in either the discovery or validation cohort” [12]). This choice of the DELFI-Pro authors means that the threshold they selected for their comparisons would be highly impacted by any outliers in the discovery cohort. Given that our methodology still obtains a specificity >99% and a PPV in excess of the 10% that is recommended for ovarian cancer screening [33], the robustness to at least one outlier appears to favor our approach. The selection of a 100% specificity threshold impacts selection of thresholds for proteins as well as the DELFI-Pro model. The >99% specificity CA125 threshold of 128.5 U/mL reported by the DELFI-Pro authors differs substantially from the 52.2 U/mL we calculate, and our threshold is much closer to the 99% threshold of 35 U/mL [5] that has been previously established within the healthy population.

With regard to ROMA, our results suggest that the ROMA approach can be successfully adapted to screening. Even without the precedent of ROMA, the combination of CA125 and HE4 makes sense to improve detection accuracy of ovarian cancer, especially in light of the clear orthogonality of the two proteins’ concentrations as seen in Supplementary Figure S1. Adding more orthogonal biomarkers to an LR model is quite reasonable, as long as the addition of the new biomarkers is shown to provide increased predictive value. As new biomarkers for ovarian cancer are investigated and shown to be orthogonal to both CA125 and HE4, it would certainly be reasonable to examine them in a combined model (be it LR or another machine learning architecture) with CA125 and HE4. It will remain important to compare such a potential improvement against a suitable baseline model, however. The evidence we presented here supports performing that comparison in a manner that guards against the influence of batch effects, and is robust to outliers in the data. Such a comparison will help to ensure that each new feature actually provides a meaningful advance, and that the complexity added by new features is merited by improvement in accuracy.

## Methods

### Data collection

We collected all data (except for library batch information) used here from the “delfipro2024” GitHub repository [34], which the DELFI-Pro authors made public as supporting material along with the DELFI-Pro publication. We also obtained in that repository scores from the protein-only screening model generated by the DELFI-Pro authors, along with the script (“code/proteins_screening_model.r”) to generate that model. Library batch data was obtained from a supplemental table of the DELFI-Pro publication, and protein values in the repository data were matched against values in another supplemental table to ensure the PGDX samples from the repository data were correctly associated with the library batches.

### Data preprocessing

As defined previously [24], the zlog transformation requires the use of a reference range derived from a healthy population, rather than calculating the mean and standard deviation (SD) from our own data. This use of a reference range is done to mitigate against outliers that could influence the mean and SD, and is based upon the 2.5th and 97.5th percentile values of a healthy population.

To perform the zlog transformation, we could not easily find firm population-wide distributions for CA125 or HE4. This led us to adopt values of 35 U/mL as the upper reference limit for CA125, and 140 pmol/L as the upper reference limit for HE4, based on previously reported highest limits [2]. For the lower reference limits, we used the recommendation [35] to set the lower reference limit to 15% of the upper limit (*i.e.*, 5.25 for CA125, and 21 for HE4), because we could not find firm guidance on a lower reference limit for either protein. These reference limits served as our estimates for the 2.5th and 97.5th percentile values for the two proteins.

### Logistic regression model training

When building our regression models, we used default values for all parameters except for one. The one exception was the class_weight=’balanced’ argument, which ensured that even though there were approximately twice as many noncancer samples as cancer samples in the discovery cohort (for either of the screening or diagnostic contexts), the class imbalance would be accounted for in the model’s fitting. This use of default values means our regularization penalty was an L2 (ridge regression) penalty term that was weighed equally with the error term of the objective function [36].

### Generation of a new feature heatmap

For ease of interpretation, we restricted our analyses to only the chromosomal copy number features and the two protein zlog values. To highlight the impact of the prefix-associated batch effect, we performed an analysis of variance (ANOVA) within the cancer samples, grouped by prefix. Features were sorted by ANOVA *p* values, with lower *p* values on the left. To make the aberrant feature values more apparent, we standardized each feature prior to plotting, subtracting the mean of each feature and dividing by each feature’s standard deviation. So that readers can see the ability to detect the batch effect on the raw data with a regular clustering, we provide a heatmap where rows and columns are clustered, and values are not standardized, as Supplementary Figure S7.

### Calculation of feature AUC values

When calculating the AUC for individual features’ ROC curves (shown in Supplementary Figure S5), we computed the AUC as a. The reported AUC was the maximum of a and 1-a, to account for the fact that some features would need to be multiplied by -1 to positively correlate with the target variable. This has the impact of making 0.5 be the minimum reportable AUC for these purposes. We used 0.75 as the threshold for declaring a feature “important” here, following previous advice [25] that features below such a threshold would not tend to be clinically useful.

### Generation of the restricted DELFI-Pro model

To create the restricted DELFI-Pro screening model, we made a copy of the DELFI-Pro authors screening model script, and made edits so that samples with the “PGDX” prefix would be removed from the metadata, training, and testing data. We made other small edits to make the script save the new model and outputs in our working directory. Additionally, we wrote code to interact with both the published DELFI-Pro model and this restricted model so that we could calculate feature importances. Scaled coefficients for the logistic regression models were calculated by dividing a feature’s coefficient by the standard deviation of that feature within the training dataset. A feature’s importance was the absolute value of its scaled coefficient divided by the sum of all features’ absolute scaled coefficients. We found this method recreated the percentages reported by the DELFI-Pro authors [12].

### Software analyses

We used the scikit-learn package [37] (v0.23.2) to build our logistic regression models and calculate classification statistics, matplotlib [38] (v3.5.1) and seaborn [39] (v0.11.2) to create plots, and pandas [40] (v1.3.5) and numpy [41] (v1.21.5) for operating on tabular data and doing general arithmetic. All code was written in Python (v3.10.12) using a Jupyter [42] (v6.4.8) notebook, with the exception of some R [43] (v4.1.2) code that we wrote to interact with the DELFI-Pro models directly.

## Data availability

All code we wrote for this project, along with copies of the data we used, are archived with and freely available from Zenodo [44]. Per-sample model scores are available within that same archive, as are tables containing the feature importances of the two DELFI-Pro screening models we compare in this work.

## Supplementary Figures

**Supplementary Figure S1.**
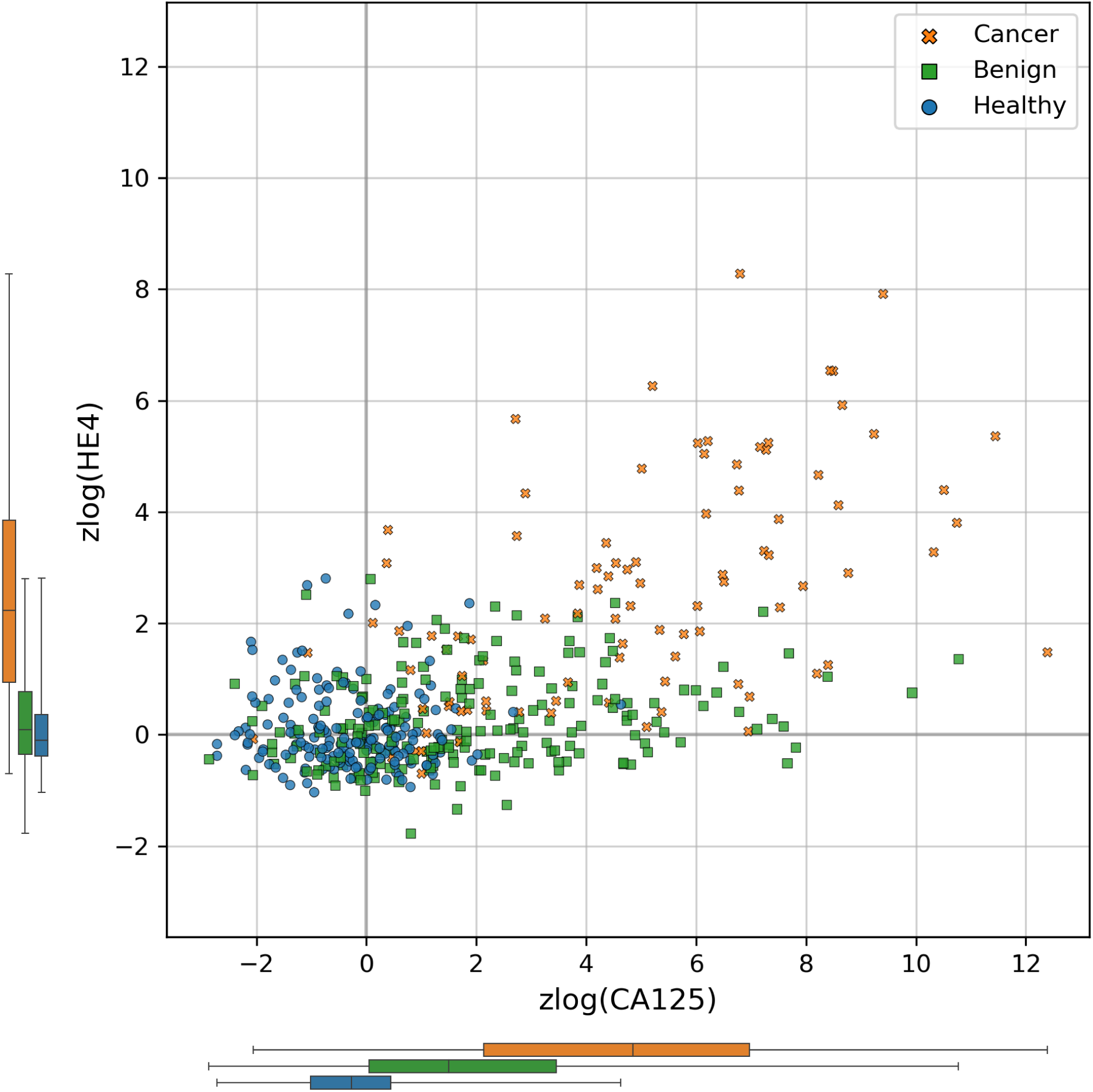
Scatter plot of zlog-transformed protein concentration values in the discovery cohort. Marginal distributions of the two protein levels within the three conditions of interest are shown to the left and under the corresponding axes, with boxplots indicating the interquartile range and whiskers representing the range from the minimum to maximum values.

**Supplementary Figure S2.**
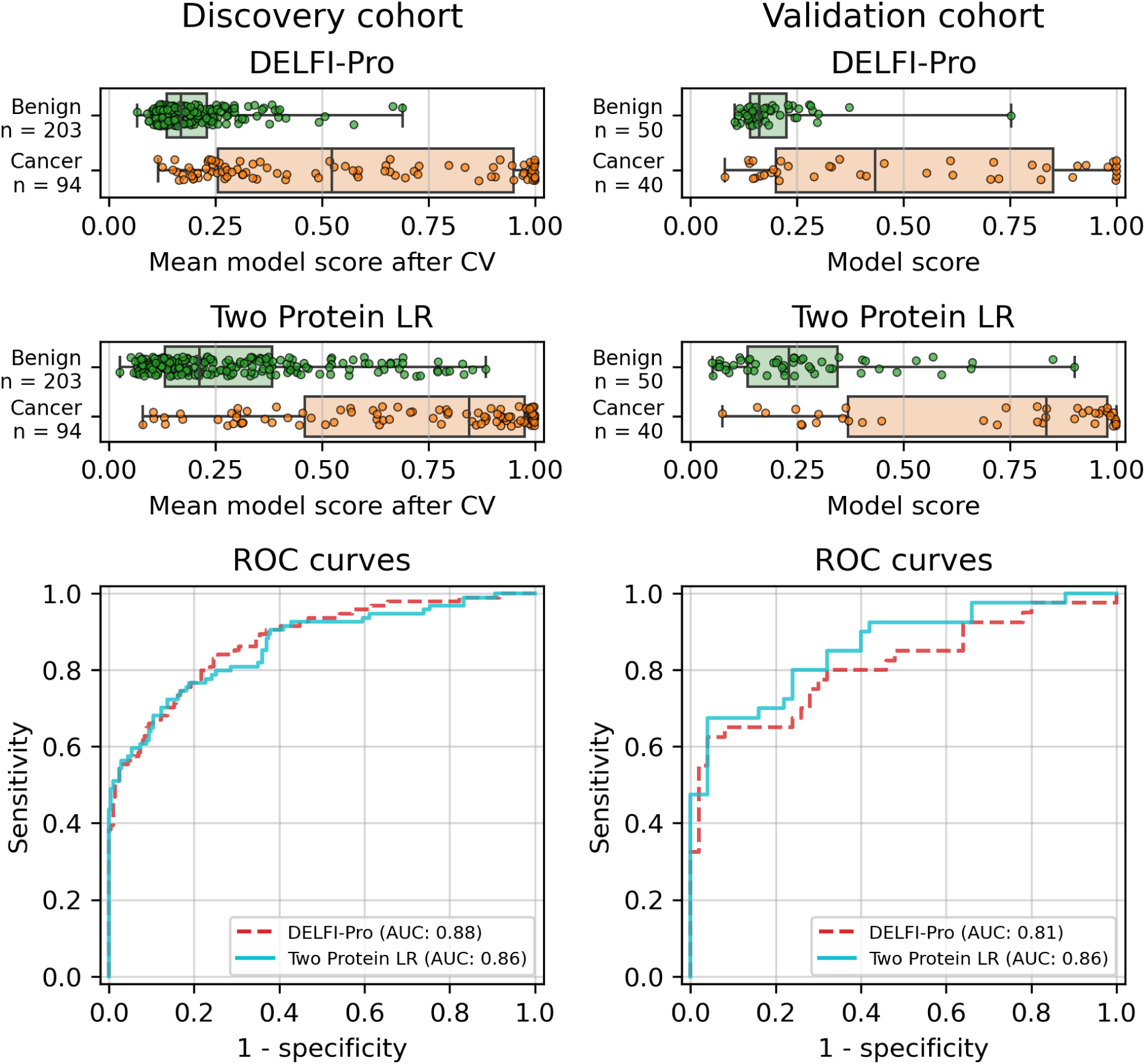
Logistic regression model performance in a diagnostic context. Cross validation (“CV”) results for the discovery cohort, and validation results on the validation cohort are shown for both the DELFI-Pro diagnostic model and a two-protein logistic regression (“Two Protein LR”) diagnostic model. Both models seek to distinguish between samples from donors with benign adnexal masses and samples from patients with ovarian cancer. Receiver operating characteristic (ROC) curves demonstrate the performance of the two models at various score thresholds, with the area under the curve (AUC) growing proportionally to model accuracy.

**Supplementary Figure S3.**
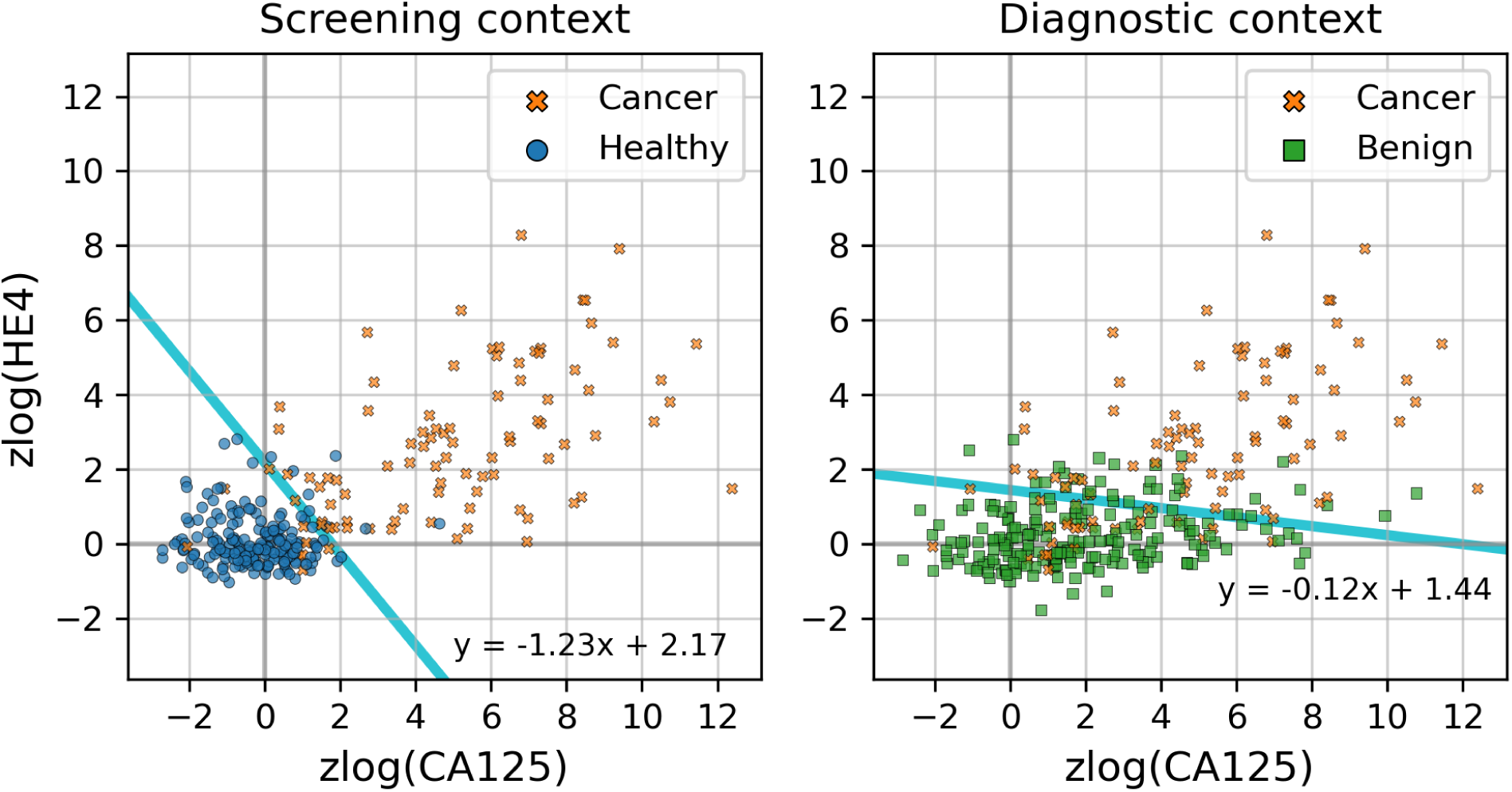
Decision boundaries learned by logistic regression models on the discovery cohort. On the discovery cohort, two protein-only models were fit to the data. In the screening context, the model distinguishes between samples from healthy donors and samples from patients with ovarian cancer. In the diagnostic context, the model distinguishes between samples from donors with benign adnexal masses and samples from patients with ovarian cancer. The decision boundaries for the two models are drawn in cyan, and equations of the decision boundary lines are written near the lines.

**Supplementary Figure S4.**
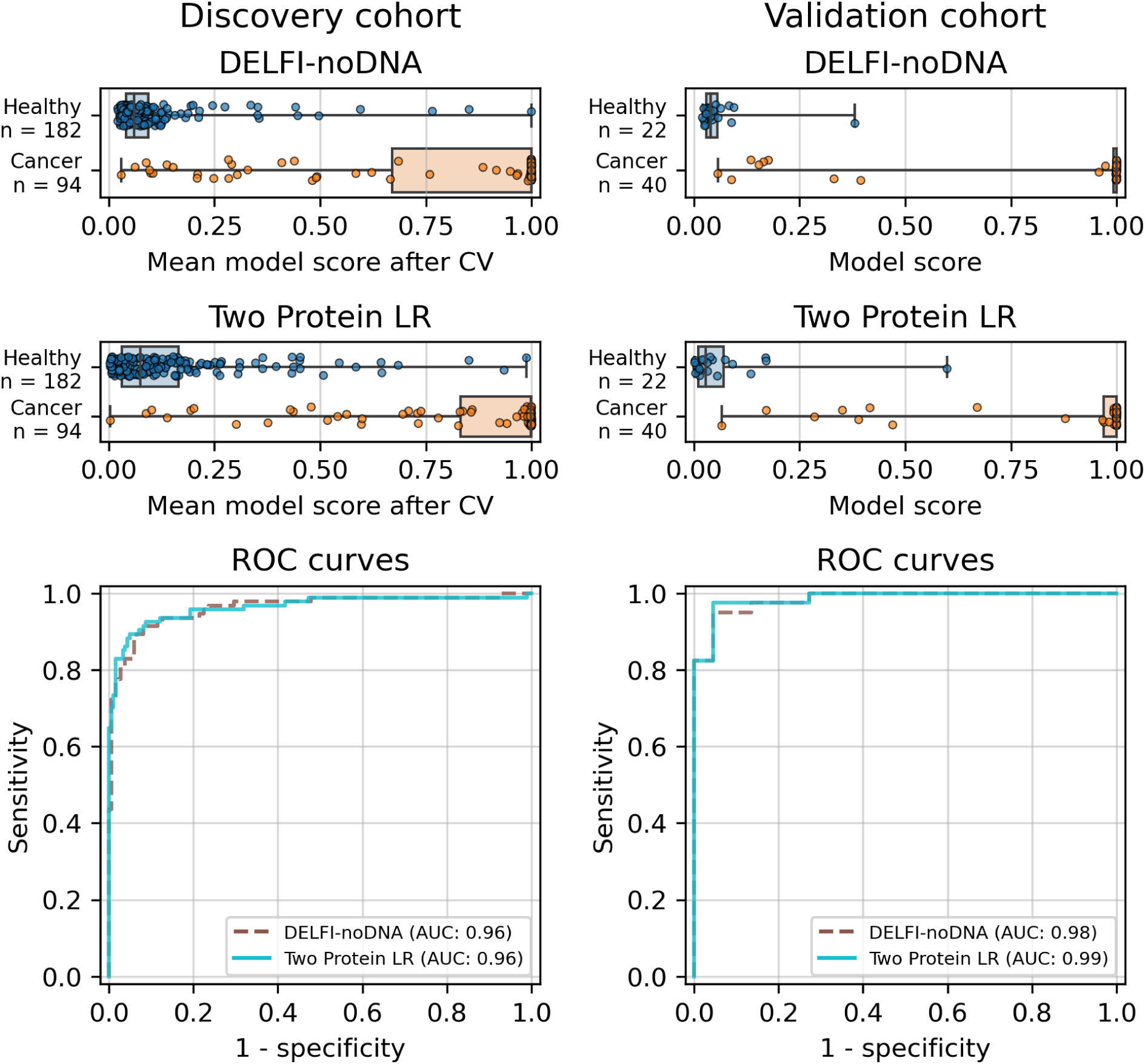
Comparison of our protein screening model with a model made available by DELFI-Pro authors. Cross validation (“CV”) results for the discovery cohort, and validation results on the validation cohort are shown for both the DELFI-Pro authors’ two-protein logistic regression screening model (“DELFI-noDNA”) and our two-protein logistic regression (“Two Protein LR”) screening model. Both models seek to distinguish between samples from healthy donors and samples from patients with ovarian cancer. Receiver operating characteristic (ROC) curves demonstrate the performance of the two models at various score thresholds, with the area under the curve (AUC) growing proportionally to model accuracy.

**Supplementary Figure S5.**
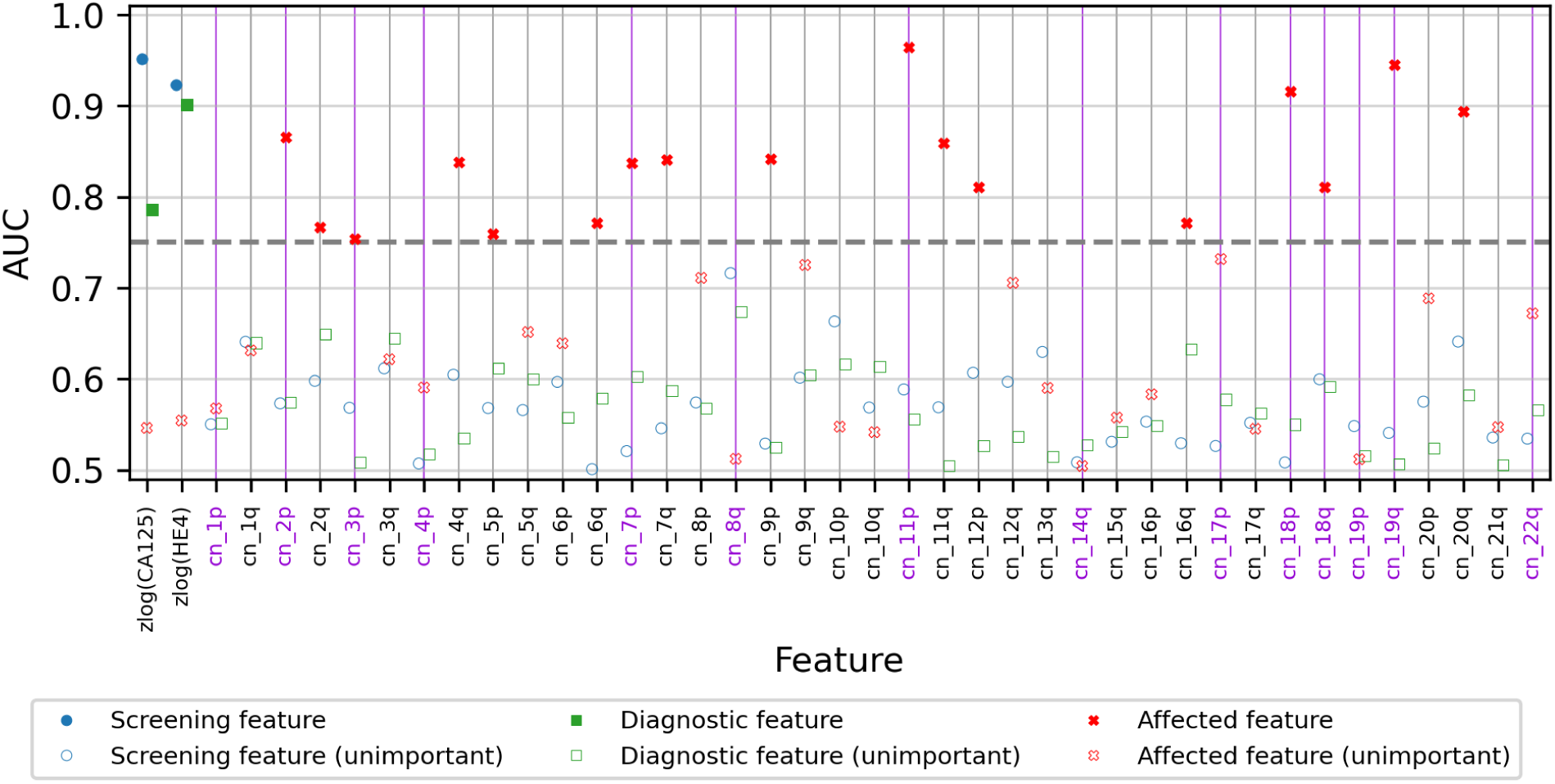
Areas under the curve for features measured in various contexts. For each displayed feature, we analyzed the discovery cohort and computed three receiver operating characteristic (ROC) curves and measured the area under the curve (AUC). Two curves were calculated on CGPL-prefixed samples, measuring a feature’s association with cancer in a screening and a diagnostic context. The third curve was calculated on cancer samples only, and measured a feature’s association with the two different prefixes (CGPL and PGDX) to determine how affected a feature was by batch confounding. Features with a “cn_” prefix are copy number features, and those with a “zlog” prefix are the zlog-transformed protein concentrations; copy number features listed in the DELFI-Pro results as contributing to the DELFI-Pro screening model performance are highlighted in violet. We use an AUC threshold of 0.75 to separate features that are good predictors from those features that are less important; features described as “unimportant” in the figure are those with AUC values less than 0.75.

**Supplementary Figure S6.**
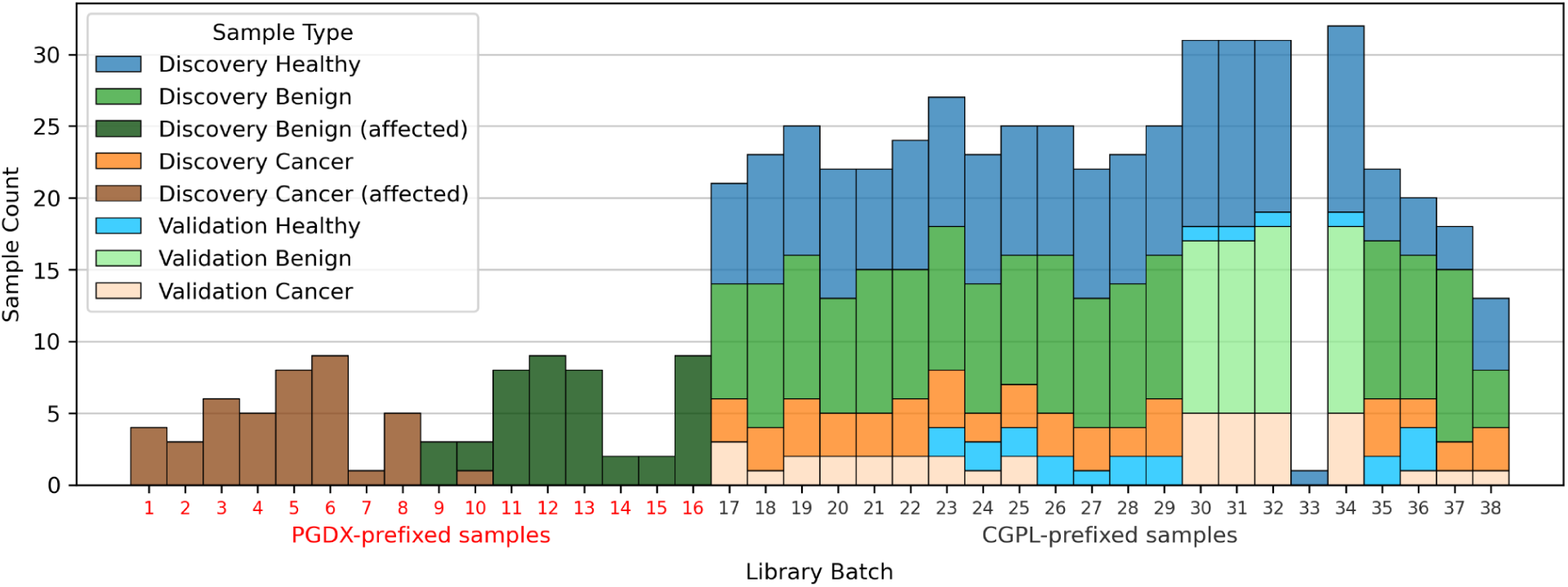
Histogram of various sample types by library batch. All samples used as part of the DELFI-Pro publication’s dataset are divided into three conditions (healthy, benign, and cancer), and into two cohorts (discovery and validation). Samples with the “PGDX” sample prefix-associated effect (labelled “affected”) are colored separately. The 16 batches that only have affected samples are colored red, with the other batches that only have CGPL-prefixed samples colored gray.

**Supplementary Figure S7.**
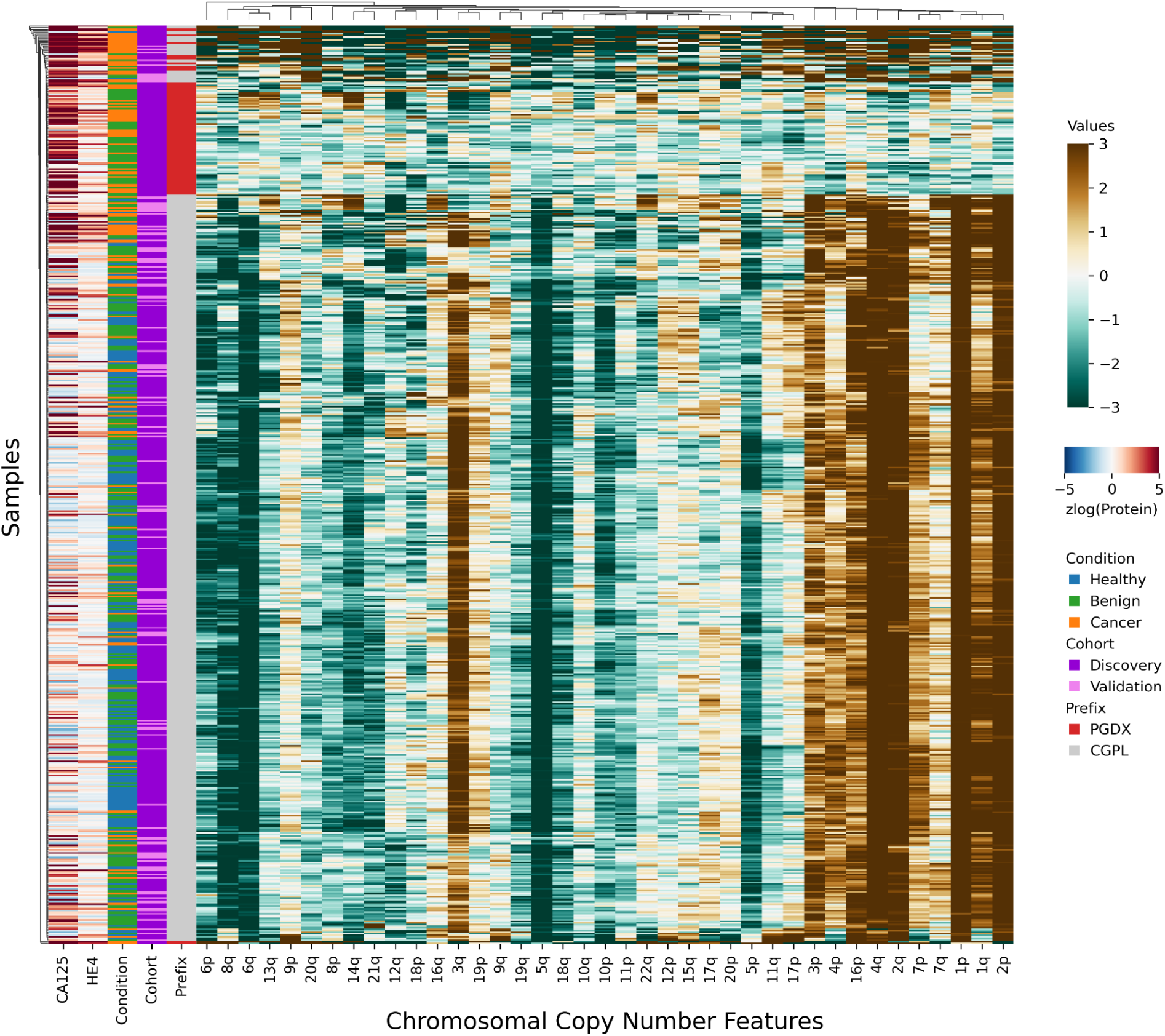
Clustermap demonstrating sample prefix-associated effect on copy number features. The samples from the discovery cohort are shown here, with each row representing features from a single sample.

